# A periplasmic metallochaperone (PmcY) couples Zn^2+^-transport to sensing in *Pseudomonas aeruginosa*

**DOI:** 10.1101/2025.05.19.654957

**Authors:** Paula Mihelj, Tomás Moreyra, Monserrat Olea-Flores, María Elena Carrizo, Teresita Padilla-Benavides, Daniel Raimunda

**Author notes:** CONICET-Universidad Nacional de Córdoba. Argentina.

## Abstract

*Pseudomonas aeruginosa* thrive to survive in harsh conditions imposed by the host. We have previously described two Zn^2+^-transporting members of the Cation Diffusion Facilitator family, YiiP and CzcD, inhibiting susceptibility to imipenem by decreasing the expression of the outer membrane porin OprD (A. Salusso and D. Raimunda, Frontiers in cellular and infection microbiology 7:84, 2017, https://doi.org/10.3389/fcimb.2017.00084). Here we provide evidence that a protein encoded in *yiiP*’s operon, PA3962, is fundamental to the coupling of these processes. Immunodetection assay indicates that PA3962 locates in membranes. Supporting a role in *oprD* regulation, a PA3962 insertional mutant has a significant increase of *oprD* expression levels and imipenem sensitivity, which is suppressed by gene complementation but not in presence of Zn^2+^, as opposed to the YiiP mutant. We identified 2 pairs of conserved acidic residues in a hydrophobic juxtamembrane domain. Metal binding specificity and stoichiometry was explored in wild-type and mutant versions of these. Zn^2+^ appears as the cognate metal of PA3962, with residues D40, D47 and D65 required for its coordination. As the periplasmic Zn^2+^-sensor CzcS regulates *oprD* expression, the interaction with it was analyzed *in vitro*. Interaction was Zn^2+^-dependent, and mutations of D47A or D65A abolished it. We propose a role for PA3962, hereafter periplasmic metallochaperone of YiiP (PmcY), in the context of Zn^2+^ signaling pathways in *P. aeruginosa*. The relay YiiP–PmcY, supplies Zn^2+^ and activates CzcS/CzcR, down-regulating the transcription of the imipenem-permeable OprD. This mechanism would allow *P. aeruginosa* to put a brake on unspecific mechanisms for micronutrient uptake, with potential xenobiotic entry, while the cytosolic Zn^2+^ quota is still sufficient.

## Introduction

*Pseudomonas aeruginosa* is a facultative anaerobic gram-negative bacteria and a prevalent opportunistic pathogen in respiratory infections of patients diagnosed with cystic fibrosis (1). It is considered an oligotrophic organism with a highly dynamic capacity for antibiotic resistance, metabolic adaptation, or phenotypical switching, which sometimes translates to fitness costs (2). Metabolic adaptations and antibiotic resistance involve optimization of Zn^2+^ usage in niches with low bioavailability (3). In the first case, the Zn^2+^-dependent (C+) to Zn^2+^-independent (C-) isoform switching of ribosomal subunits, increases the intracellular Zn^2+^ pool in this condition (4). In the second, Zn^2+^ induces carbapenem resistance by repressing the transcription, and then the antibiotic influx, of the outer membrane porin Occ1, also known as OprD (5). As carbapenem-resistant OprD mutants show enhanced fitness *in vivo* (6) we hypothesize that this “master of survival”, could benefit from a last resort-type of intracellular Zn^2+^ availability by finely tuning Zn^2+^-sensing mechanisms. In this way this opportunistic pathogen could adapt metabolically and become resistant to carbapenems, avoiding the pay-offs of a fitness cost imposed by nutritional immunity(7).

We have previously shown that the Zn^2+^ exporters of the Cation Diffusion Facilitator family, YiiP and CzcD regulate levels of OprD (8). Single insertion mutants of these genes showed increased sensitivity to imipenem, and increased Zn^2+^ sensitivity when introduced into double mutant backgrounds where other Zn^2+^ resistance mechanisms were also affected. Considering studies by others showing that transcription of *czcD* responds to Zn^2+^ levels in the growth media with a slow temporal activation (9) a role in Zn^2+^ detoxification might be considered. However, the fact that *yiiP* transcripts are not upregulated by Zn^2+^, opens new possibilities for the role of this Zn^2+^-transporter in Zn^2+^ mediated signaling.

Zn^2+^ sensing mechanisms in *P. aeruginosa* are well known. At the cytoplasmic level, CadR, a member of the MerR transcription factor family, controls the expression of CadA, also known as ZntA, a Zn^2+^-transporting P_IB_-ATPase (8, 10). Interestingly, it has been proposed that under high Zn^2+^ conditions, CadA is required for timely expression of CzcCBA, also known as HME-RND, a tripartite Zn^2+^-export system that promotes efflux from the periplasm to the extracellular space (10). This implies that the specific CadA Zn^2+^-export is coupled to Zn^2+^ sensing in the periplasm. In line with this, extracellular Zn^2+^ bioavailability in *P. aeruginosa* is primarily detected in the periplasm via the two-component system (TCS) CzcS-CzcR. CzcS senses periplasmic Zn^2+^ by coordinating the metal in the periplasmic domain (11), which is transduced to cytosolic domains promoting autophosphorylation of the histidine kinase domain and finally, the transphosphorylation to CzcR. Phosphorylated CzcR (P-CzcR) has dual effects on transcription of genes in the CzcS-CzcR regulon (12). For example, P-CzcR up-regulates transcription of HME-RND, thus counteracting a potential Zn^2+^ overload (9, 10) in the periplasm. However, P-CzcR affects negatively the transcription of genes involved in pyocyanin synthesis and of the outer membrane porin *oprD* (11, 12).

Here we present evidence suggesting that a <u>p</u>eriplas<u>mic</u> metallochaperone of <u>Y</u>iiP (PmcY/PA3962) is required for *oprD* repression and thus, for Zn^2+^ sensing in the periplasm. Our data shows that PmcY is a Zn^2+^-metallochaperone that interacts with CzcS, its putative acceptor or target protein. We also propose that the intracellular Zn^2+^ efflux to the periplasm mediated by YiiP-PmcY has coevolved in Pseudomonadales as a mechanism for limiting the fitness cost of adapting to common environmental or host-imposed scenarios with low Zn^2+^ bioavailability, by repressing an unspecific outer membrane porin.

## Materials and methods

### Bacterial strains, plasmids and growth media conditions

*Pseudomonas aeruginosa* MPAO1 wild-type (WT), and the insertion mutants *yiiP::dTn5* and *pmcY*/PA3962*::dTn5* were obtained from the *P. aeruginosa* Mutant Library at the University of Washington ((13); Table 1). For complementation of the *pmcY*/PA3962*::dTn5* strain PA3962 locus plus the 250bp upstream bases was PCR amplified and cloned in pSEVA621 using EcoRI and KpnI sites (14). The sequence for *gfp* was introduced at the 3’-end as a fusion in order to promote protein stability using KpnI and XbaI sites. Electrocompetent cells were transformed by electroporation as described (15) and selected on Luria–Bertani (LB) agar plates supplemented with gentamicin (30 µg/ml). Double mutant *oprD-pmcY* (DM3004) was obtained via double recombination events following methods previously described (16). The up and down-stream 500 bp fragments of PA3962 were amplified and stiched by PCR. After purification and digestions with EcoRI and HindIII restriction enzymes the fragment was ligated into the suicide plasmid pEX18-Gm. Electrocompetents *oprD::dTn5* cells were transformed with this plasmid and recombinants were selected on LB-agar supplemented with 60 µg/ml gentamicin. Several colonies were grown in LB supplemented with sucrose 5%. The PA3962 locus deletion was confirmed by PCR using genomic DNA of isolated colonies grown in LB + sucrose 5%, supplemented with tetracycline (30 µg/ml). The primers used in these and other procedures are listed in Table 1.

**Table 1.**
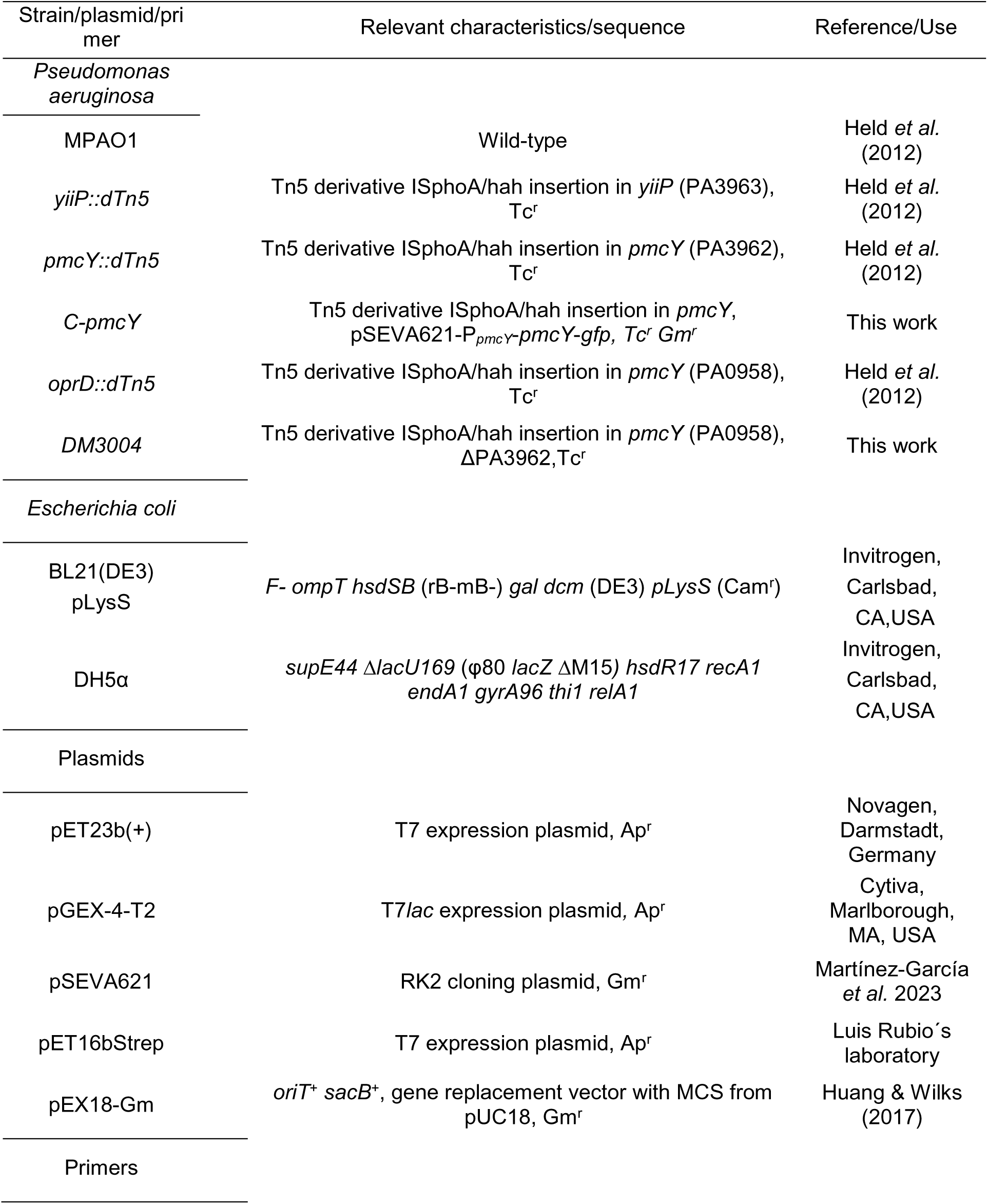
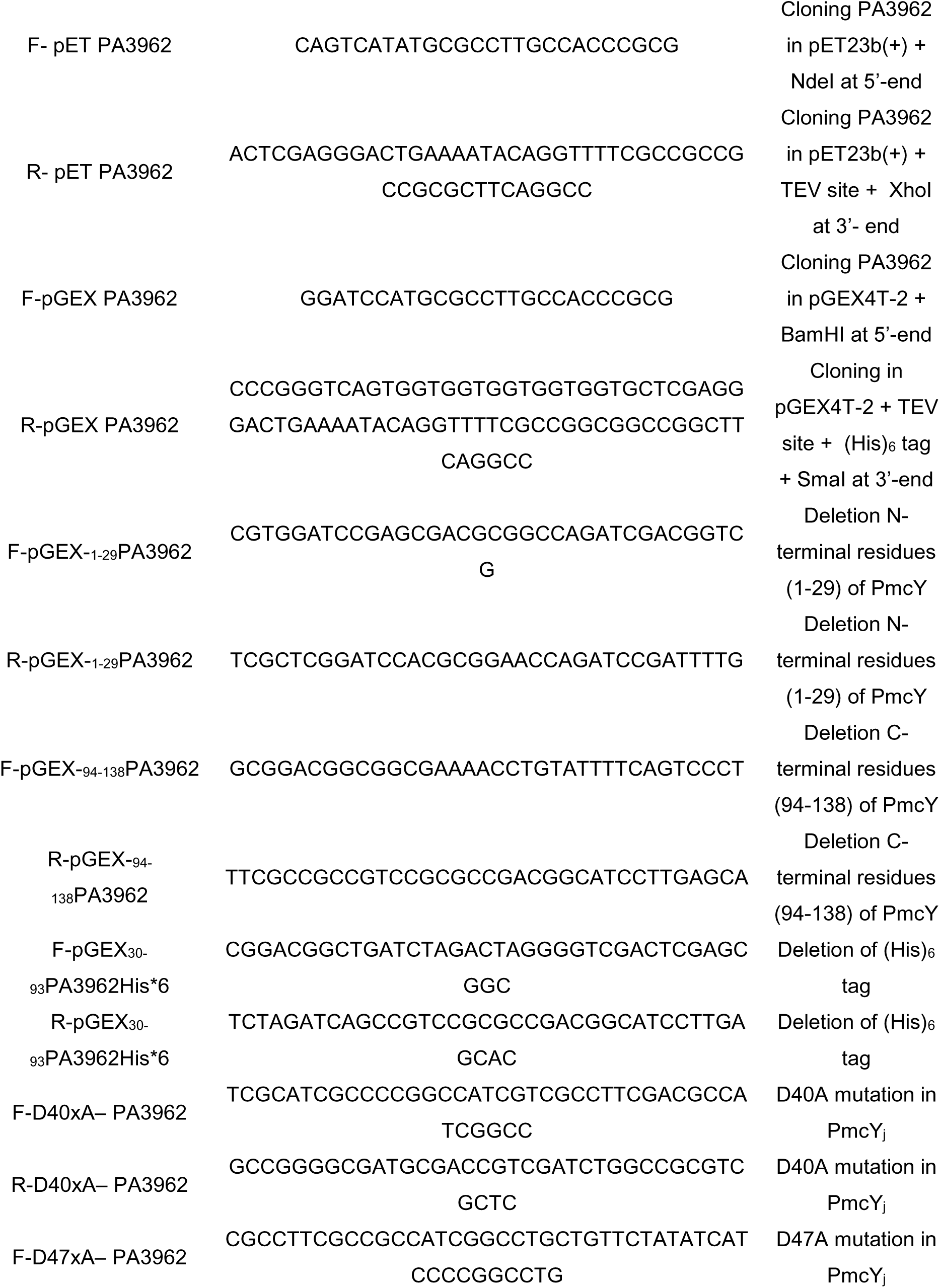
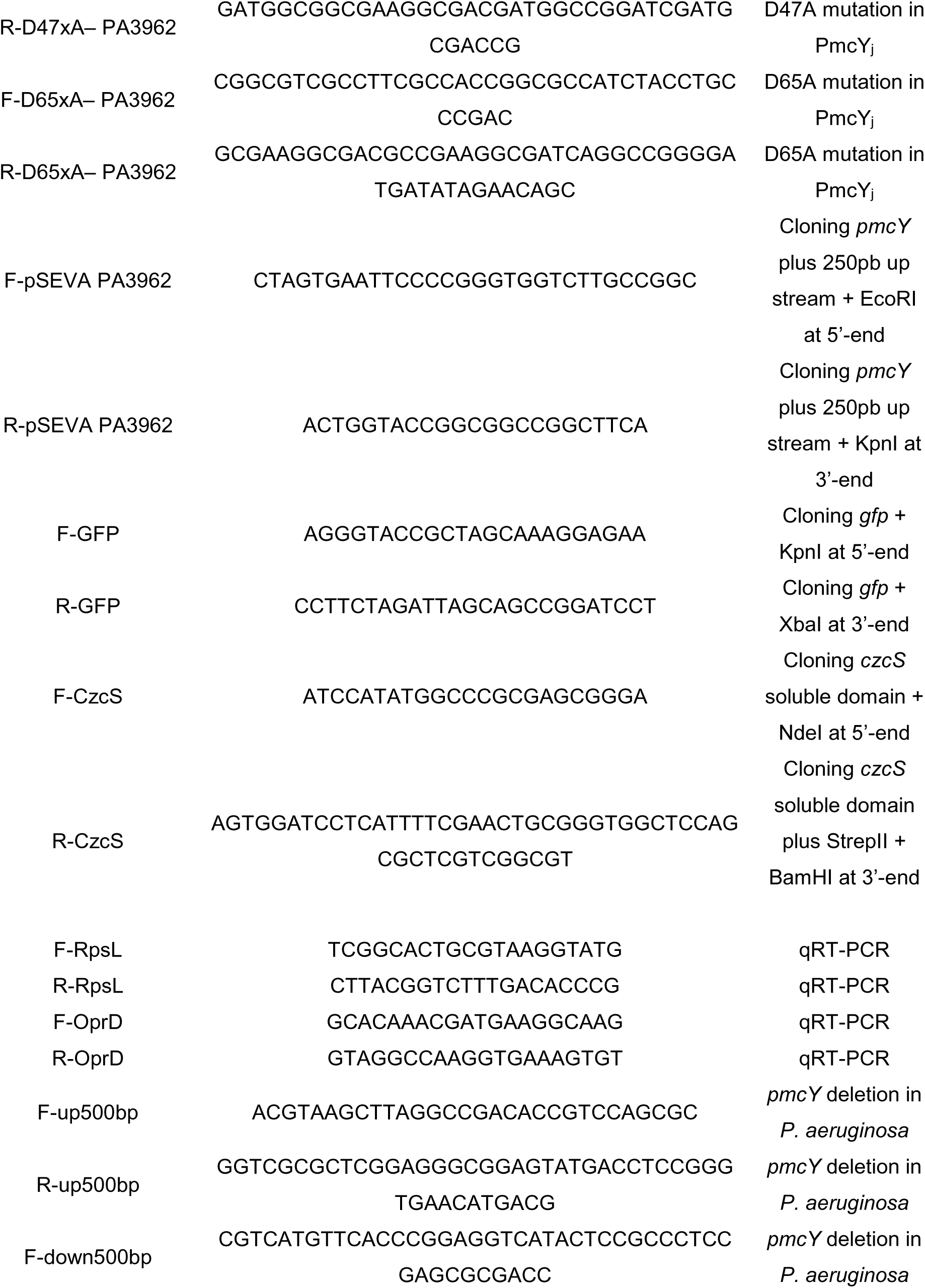
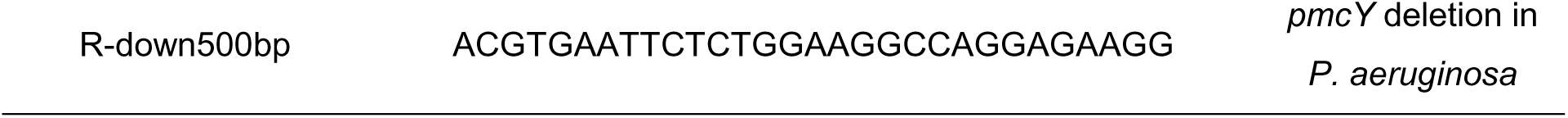
List of Strains, plasmids and primers used in this study.

### Plasmid constructions and protein expression and purification

In a first attempt to yield a whole version of PmcY by heterologous expression in *E. coli* PA3962 locus was cloned from *P. aeruginosa* MPAO1 gDNA and inserted by ligation in the expression plasmids pET23b(+) (Novagen) or pGEX4T-2 using primers described in Table 1. These constructs contained at C-terminal a hexahistidine ((His)_6_) tag preceded by Tobacco Etch Virus (TEV) protease recognition site. Due to low yield of the protein and guided by bioinformatics analyses we expressed the juxtamembrane domain of PmcY (PmcY_j_) as a GST fusion protein by deletion of N- and C-termini and (His)_6_ tag of PmcY by the one-step site-directed deletion PCR protocol (17). Site directed mutagenesis of conserved residues were obtained using the same procedure. Similarly, the soluble domain (SD) of CzcS (amino acids 40-166, CzcS_SD_) was cloned by PCR amplification from MPAO1 gDNA and ligated in pET16bStrep (Table 1) with a twin-strep tag at N-terminal. All constructs and mutants were confirmed by DNA sequencing.

For protein expression and purification *E. coli* BL21(DE3)pLysS were transformed with one of the above constructs. After selection cells were grown in LB overnight at 37°C on a shaker at 200 rpm. Next day 1-2 L of LB media with antibiotic was inoculated with cells to reach OD_600_= 0.1 and grown in the same conditions till early log-phase. At this point cultures were induced with 0.15 mM IPTG for 16 h at 20°C. For purification of GST and GST fusion versions of PmcY_j_ cells were resuspendend in 50 ml of Tris-HCl 20 mM pH 7.5, NaCl 500 mM, TCEP 0.25 mM, 0.1% Triton X-100 (Buffer A) plus PMSF 1 mM, and lysed in a cell disruptor EmulsiFlex C3 (Avestin). After ultracentrifugation at 100,000 *g* the supernatant was incubated with 1 ml of Glutathione Sepharose 4B or Fast Flow (GE Healthcare) for 1 h at room temperature. Resin was packed by gravity in a polypropylene column (Bio-Rad) and washed with 30 resin volumes of buffer A. The fusion protein was eluted with buffer A containing 40 mM glutathione. The soluble domain for CzcS was expressed and purified as described earlier (11) with minor modifications. After growth and induction cells were lysed as above and ultracentrifuged at 100,000 *g*. The supernatant was incubated for 1 h at room temperature with 0.5 ml of Strep-Tactin resin (IBA Lifesciences) equilibrated with buffer A for 1 h and eluted after washing with 30 resin volumes. Bound protein was eluted with 10 resin volumes of buffer A containing 50 mM of D-biotin. In both cases elution fractions were analyzed by SDS-PAGE 12%, Coomassie brilliant blue staining and Western blot using (His)_6_-tag antibody (pAb, Rabbit, Genscript), GST-tag antibody (pAb, Rabbit, Genscript) or StrepMAB-Classic antibody (IBA Lifesciences) as primary antibodies. Those fractions containing protein were pooled for dialysis in buffer A using a dialysis tubing (SnakeSkin, Thermo Scientific) with 3.5 kDa cut-off. After addition of 10% glycerol, protein concentration was estimated by Bradford (Bio-Rad) and 280 nm lectures in NanoDrop (Thermo Scientific). Protein samples were frozen and maintained at - 80°C for later use. For in-column interaction assays the bait protein, GST-PmcY_j_ WT and mutant versions, were not eluted from the resin and kept at 4°C in the dark for later use and store for no more than 2 weeks. In this case protein concentrations were determined by densitometry. Denatured resin samples were run in SDS-PAGE 12%, stained with Coomassie brilliant blue (CBB) and images analyzed with ImageJ (18). For calibration, bovine serum albumin (Sigma-Aldrich) standards were run in the same gel.

### Antibiotic sensitivity assays

Susceptibility to imipenem (Sigma-Aldrich) and ciprofloxacin was assessed by the agar dilution or the inhibition halo methods in Müller–Hinton (MH) medium as described before (8). For agar dilution overnight grown cells were serially diluted from an initial OD_600_= 1 and then 10 µl were spotted on MH agar plates with different antibiotic concentrations. For inhibition halos, a suspension of cells at OD_600_=0.05 in soft agar were layered on top of MH agar plates. After solidifying, 7-mm diameter filter paper discs embedded with the indicated amount of imipenem were located on top of the soft agar. In all cases growth or inhibition was evaluated after overnight incubation at 37°C. Analyses of the imipenem MIC values found by agar dilution method (ADM) *vs*. our previously data published using Epsilon-test method (Biomerieux Diagnostics) yielded a lineal correlation between MICs following *y=8.082 x* with and R² = 0.797, where *y* equals the apparent ADMs MICs, and *x* the Epsilon-test ones. Thus, apparent imipenem concentrations are informed in this work.

### Quantitative RT-PCR analysis

Overnight cultures from *P. aeruginosa* WT, *yiiP::dTn5* and *pmcY::dTn5* were diluted to an OD_600_ of 0.05 in LB and incubated until they reached an OD_600_ of 0.6-0.8. Total RNA was extracted with TRIzol^TM^ Reagent (Invitrogen) according to the supplier’s instructions and treated with RNase-free DNase I (TransGen Biotech) for 2 h at 37°C to remove residual DNA. After phenol/chloroform extraction, RNAs were precipitated with ethanol and the pellets resuspended in RNase-free water. For cDNA synthesis, 200-300 ng of RNA was reverse transcribed using random primers and *EasyScript*^®^ Reverse Transcriptase (TransGen Biotech) according to the manufacturer’s instructions. Quantitative PCR was performed using FastStart Universal SYBR Green Mastermix (Rox) (Roche), according to the supplier’s instructions. Fold change expression levels were calculated using the 2^-ΔΔCt^ method as previously described (19). Results were normalized with the *rpsL* gene (PA4268). Primers used for qPCR studies are listed in Table 1.

### Metal binding assays

Transition metal (TM) binding capacity of GST fusion proteins was evaluated using the chromogenic dye 4-(2-Pyridylazo) resorcinol (PAR, Tokyo Chemical Industry). For incubation of the proteins with TM we followed described protocols for in-solution (20) or in-column (21) assays with some modifications. To assess binding to proteins 10-20 µM samples (or the mass bound to resin to yield this concentration before adding PAR) were incubated for 30 min with 5 molar excess of TM in buffer A. For in-solution binding assays unbound TM were separated by exclusion chromatography with Sephadex G25-80 (Sigma-Aldrich) and for the in-column type the resin was washed with 10 volumes of buffer A to remove unbound TM. Before TM quantification protein concentrations were estimated as described earlier and samples containing proteins in solution or bound to resin were treated with 0.4-0.8 mg/ml of proteinase K. For TM quantification, PAR was added to a final concentration of 100 µM and read in microplates at 500 nm. Calibration curves were done in parallel for each TM and the molar ratio of protein to TM calculated. In both conformations GST alone or GST fusions were run in parallel to detect TM bound in proteins as purified. These values represented no more than 10% of those found in TM incubated samples and were subtracted in all cases. For the quantification of Mn^2+^ the buffer pH was modified from 7.5 to 11 as the colorimetric reaction of this TM is more sensitive. Therefore, after the digestion reaction with proteinase K, the samples were evaporated completely and resuspended in 200 mM CAPS (3-(Cyclohexylamino)-1-propanesulfonic acid) buffer pH 11.

### Protein localization

The *E. coli* strain BL21(DE3)pLysS transformed with the plasmid pET-23b(+)-PA3962, which expresses the PA3962 protein fused to a 6-histidine tag, was grown in 12.5 ml of LB medium at 37°C for 16 h. This culture was placed in two flasks with 500 ml of LB medium and grown with shaking (200 rpm) at 37°C for 1 h. One of the cultures was induced with 0.15 mM of IPTG and the other was used as a control (basal). After 3 h of growth the cells were harvested by centrifugation at 4,000 *g*, 4°C for 10 min. Cell pellets were washed in ice-cold 30 ml of 50 mM Tris-HCl pH 7.5 and 0.3 M sorbitol (buffer S) 3 times, with addition of 10 µM EDTA in the second wash.

Cells were pelleted by centrifugation at 9,000 *g*, 4°C for 10 min. Finally, cells were resuspended in 30 ml of ultrapure water and incubated for 5 min at room temperature. They were then centrifuged at 11,000 *g*, 4°C for 30 min. The supernatants were collected and centrifuged again to eliminate any possible contamination. After this they were considered the periplasmic fractions. For cytosolic and total membrane fractions cell pellets were resuspended in 7 ml of 10 mM Tris-HCl pH 7.5 and disrupted at 4°C using a tip sonicator (Vibra Cell, Sonics & Materials, Inc.) with 6 cycles of 20 seconds on and 60 seconds off pulses. The lysate was centrifuged at 6,000 *g* for 10 min at 4°C to remove undisrupted cells and centrifuged again at 100,000 *g* for 30 min at 4°C. The supernatants were recovered and classified as cytoplasmic fractions. The resultant pellets were resuspended in 5 ml of 10 mM Tris-HCl pH 7.5 and classified as total membrane fractions. The protein concentration of all fractions was quantified by Bradford assay, and those with a value less than 1 mg/ml were concentrated using centrifuge filtration tubes (Centricon, Millipore). Samples of all fractions were prepared at concentrations of 1 mg/ml and 5X Laemmli sample buffer was added. A final total protein mass of 10 µg was loaded per lane on 14% SDS-PAGE gels in duplicate, one for CBB staining and the other for Western blot. (His)_6_-tag antibody (pAb, Rabbit, Genscript) and anti-rabbit antibody labeled with horseradish peroxidase were used to detect expression in fractions. Images were taken using Amersham Image Quant 800 Fluor (Cytiva).

### Protein-protein interaction assay

To evaluate the interaction between PmcY and CzcS, the recombinant proteins GST-PmcY_j_ (WT and mutants D40A, D47A, and D65A) and CzcS_SD_ were used. A 100±20 mg sample of purified WT or mutant versions of GST-PmcY_j_ were treated with EDTA-Tris. For this, Glutathione Fast Flow (GE Healthcare) resin with bound protein was incubated with 1 ml of buffer A plus 1 mM EDTA-Tris in a 10 ml polypropylene chromatography column (Bio-Rad). Three ml of buffer A were used to remove residual EDTA from column. The interaction was tested in two conditions, with GST-PmcY_j_ loaded with (+Zn^2+^) and without (-Zn^2+^) ZnCl_2_. For this, resin was incubated for 5 min with 500 µl of 100 µM ZnCl_2_ in buffer A or just buffer A, respectively. The CzcS_SD_ protein was used in solution and previously treated with 0.5 mM EDTA-Tris for 30 min in ice. EDTA-Tris was removed with PD-10 column equilibrated in buffer A. In-column GST-PmcY_j_ and mutant versions, in its two conditions (+Zn^2+^) and (-Zn^2+^), were incubated with 25 μM CzcS_SD_, added in 250 μl, during two passes through the column (Flow through, FT) and 4 washes (W1-4) were performed with 250 μl of buffer A. Finally, the resin was resuspended in 250 μl of buffer A and samples prepared with 5X Laemmli loading buffer for SDS-PAGE 12% analyses. After CBB staining images were taken using Amersham Image Quant 800 Fluor (Cytiva).

### Bioinformatics analyses

To analyze the conservation of residues in a putative PmcY family, homologs were searched in a local database of all protein sequences of RefSeq reference and representative genomes protein sequences (Refseq_select_prot) (January 2024) by the reciprocal best hit (RBH) method (19). Four hundred and sevenunique organisms having PmcY-like members with unique accession numbers were found. Multiple sequence alignment (MSA) was done in Clustal Omega (22) using a similar number of sequences and the same MSA was used in ChimeraX (23) to create the amino acids conservation map in the putative 3D AF structure of PA3962 (AF-Q9HX54-F1).

This number of organisms were used later to compare and create a list of non-redundant bacterial organisms having, or not, the *pmcY-yiiP* architecture. The search of the *pmcY-yiiP* operon was done using Multigenblast (24). Similar RBH searches were performed for OprD and CzcS homologs in the same data base. After removal of duplicated organism names, a 4-group diagram was prepared with these organisms (https://bioinformatics.psb.ugent.be/webtools/Venn/) and a phylogenetic tree was constructed for visualization of coevolution in iTOL (25).

## Results

### PA3962 is a membrane protein

PA3962 is part of a putative operon with YiiP (PA3963). Previous bioinformatics studies of identified *P. aeruginosa* secreted proteins pointed that the N-terminal sequence in PA3962 is a type II signal peptide (26). These results were supported with a plasmid library screening in *E. coli* where several PAO1 genomic DNA fragments, including PA3962, fused to truncated alkaline phosphatase (PhoA) drove its periplasmic localization. Signal peptides are recognized by different secretion systems in bacteria and classified in different groups. The type II are related to proteins anchored to the membrane, also called lipoproteins. A consensus sequence called “lipobox” is found in the type II signal peptide, which allows for molecular recognition by the machinery involved in processing and finally anchoring the protein to the inner or outer membrane. For this, first a conserved cysteine residue in the lipobox is diacylated with diacylgycerol. Second, the signal peptide is removed by a specific protease leaving free and reactive the amino group of the cysteine for a final third acylation (27). Figure 1, shows the predicted structure of PA3962 by AlphaFold2 (AF2) (28) showcasing the colour–coded hydrophobic profile generated in ChimeraX and position of Cys23 in the lipobox. The ribbon structures depict the putative position of the protein facing the periplasmic space after the anchoring occurs. In order to confirm the membrane localization of PA3962 we expressed the whole protein with a C-terminal (His)_6_ tag in *E. coli* in basal and induced conditions. After fractioning cells, cytosolic, total membranes, and periplasmic samples were analyzed by SDS-PAGE and Western blot using anti-(His)_6_ as primary antibody (Fig. 1B). Protein detection was evident in most of the fractions and signal was increased under basal conditions, pointing to an inhibitory effect of the translation in presence of the inducer when the full length protein is overexpressed. Most importantly the detection in total membrane fraction was observed. As the mass of protein loaded in all lanes was similar, and assuming an equal contribution of each compartment in cell mass, a qualitative conclusion is that PA3962 is a membrane protein. PhoA detection in *E. coli* requires folding in the periplasm, thus this orientation is the most likely scenario in *P. aeruginosa* and we renamed PA3962 as <u>P</u>eriplas<u>mic</u> metallochaperone of <u>Y</u>iiP, PmcY.

**Figure 1.**
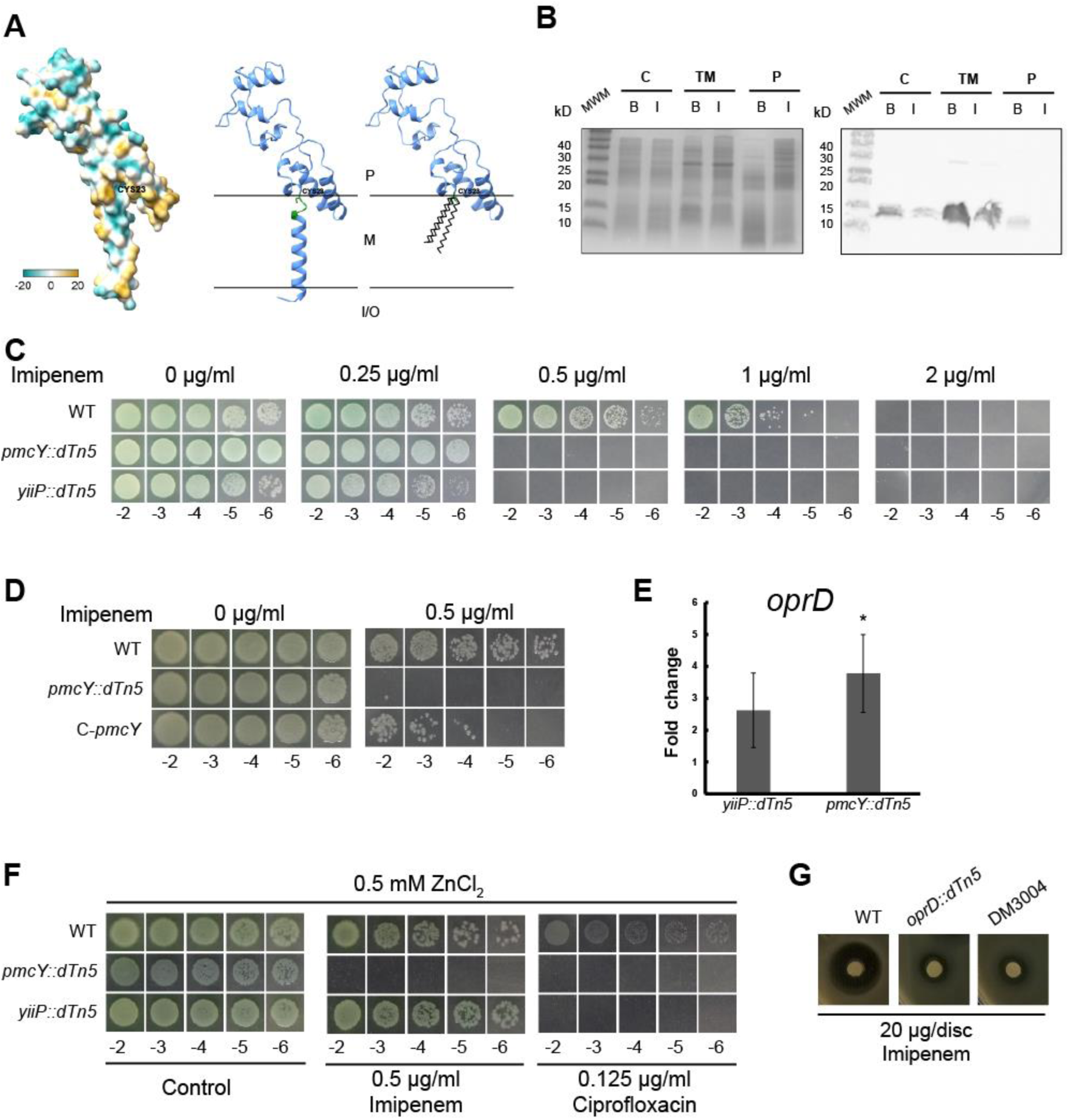
PmcY is a membrane protein that counteracts imipenem susceptibility. (A) Left, hydrophobic surface profile of the AF2 predicted structure for PmcY/PA3962. Position of Cys23 is indicated. Center and right, ribbon diagram of PmcY with lipobox amino acids indicated in green. The thiol group of Cys23 would promote acylation anchoring the protein, right. Notice that independent of the position at the inner or the outer membrane, PmcY faces the periplasmic space. P= periplasm; M= membrane; I/O= inside/outside the cell. (B) Analysis of heterologously expressed PmcY-(His)_6_ in *E. coli* compartments (cytosolic= C; total membranes= TM; periplasm= P) by cell fractioning. Ten µg of protein from each fraction obtained from cells grown in LB media (basal= B; induced = I) were analyzed by SDS-PAGE (left) and Western blot (right). PmcY-(His)_6_ expression is apparently increased in basal conditions and mostly found in the membrane fraction. (C) Serial dilutions (10 μl) of *P. aeruginosa* PAO1 (WT), *pmcY::dTn5* and *yiiP::dTn5* cells initially at OD_600nm_=1.0 were spotted on MH-agar supplemented with the indicated imipenem concentration and incubated for 16 h at 37°C. (D) Complementation of *pmcY::dTn5* restores imipenem susceptibility. PA3962 plus 250bp upstream bases were cloned in the low copy plasmid pSEVA621 as a fusion to GFP. This construct was introduced by electroporation into *pmcY::dTn5*. No complementation was observed in *pmcY::dTn5* cells transformed with empty pSEVA621 nor with pSEVA621-GFP (not shown) (E) Fold change expression levels (2^-ΔΔCt^ with *rpsL* as reference gene) of *oprD* transcripts *vs.* WT. Cells were grown until mid-log phase at 37°C in LB before RNA extraction. Data are the mean ± SE of three independent experiments. *, *P*<0.05 from a two-tailed Student’s t test of averaged ΔCt (*vs.* WT). (F) Zn^2+^ induced lack of susceptibility response is specific to imipenem and is PmcY dependent. Notice that *pmcY::dTn5* is unresponsive to Zn^2+^ and the lack of response to ciprofloxacin in presence of the TM in both mutant strains. Agar dilution method was used to determine MICs. (G) Double mutant *oprD-pmcY* (DM3004) shows similar sensitivity to imipenem as the single mutant *oprD::dTn5*. Inhibition halos were photographed after 16 h incubation at 37°C. Similar sensitivities between mutant strains were obtained with increasing imipenem masses in the filter discs (not shown).

### PmcY is involved in *oprD* mediated imipenem susceptibility

As previous results indicated a role for YiiP in Zn^2+^-dependent antibiotic susceptibility involving OprD and altered membrane permeability (8) we next analyzed if absence of PmcY yielded similar effects. The minimal inhibitory concentration (MIC) for the antibiotic in Tn5-generated mutant strains (Fig. 1C) YiiP and PmcY mutants showed that both are sensitive to imipenem *vs.* WT. Complementation of PmcY strain was achieved by transformation of mutant cells with the low copy plasmid pSEVA621 (14) carrying PA3962 locus under transcriptional regulation of 250 bp upstream sequence (Fig. 1D). We have previously demonstrated that lack of YiiP induces expression of OprD (8). If PmcY is involved in imipenem susceptibility the expression of *oprD* should be altered compared to the WT strain. The results of *oprD* locus expression between PmcY and YiiP mutant strains (*vs*. WT) correlates to the imipenem susceptible phenotype of both (Fig 1E). These results indicated that the molecular mechanism leading to the loss of imipenem susceptibility involves the lack of Zn^2+^ sensing capacity. This loss would involve specifically Zn^2+^ transport from cytosol to the periplasm. Thus, we hypothesized that both, YiiP and PmcY, mutants would remain sensitive to imipenem at high Zn^2+^ content (0.5 mM) in the growth media. However, only PmcY mutant strain showed the same imipenem susceptibility in presence of high levels of Zn^2+^ indicating a key role for PmcY. The apparent dispense of YiiP to sense Zn^2+^ at high extracellular Zn^2+^ conditions, correlates with a save-for-later strategy to sustain steady-state levels of the intracellular quota of this vital element. To test the Zn^2+^-specificity of the mechanism we tested also ciprofloxacin sensitivity (Fig. 1F right). The lack of Zn^2+^ complementation to this antibiotic susceptibility observed in both mutants, points to the involvement of another mechanism likely related to membrane permeability of ciprofloxacin and other more lipophilic antibiotics, as suggested previously (8). To discard any agonistic effects between OprD and PmcY related to membrane permeability, we generated the double mutant *oprD-pmcY* (DM3004) and evaluated its imipenem susceptibility. Figure 1G shows that both strains, *oprD::Tn5* and DM3004, behave similarly. These results indicate that PmcY is required for Zn^2+^ sensing in the periplasm and that *oprD* is regulated by PmcY activity. Thus we focused on the pivotal role of PmcY as part of a sensing mechanism leading to decrease intrinsic imipenem susceptibility, and hypothesize that PmcY should bind Zn^2+^ and interact with a sensor to likely transfer the metal to it.

### PmcY binds Zn^2+^ specifically through conserved aspartate residues

We hypothesized that PmcY plays a role as metallochaperone. These function in metalation of proteins or in detoxification pathways, and in both cases the commonality is their capacity to bind a specific transition metal through conserved residues. A multiple sequence alignment of PmcY homologs obtained by a reciprocal best hit search in complete bacterial genomes allowed us to identified conserved residues (Fig. 2A, maroon). Most of these would be located in a periplasmic domain juxtaposed to the membrane. Conserved amino acids are all acidic, with oxygen (O) ligands of lateral chains of aspartates and glutamates. After several failed trials of full length PmcY expression and purification to get enough protein for biochemical characterization, we opted to fuse this domain (amino acids 30 to 93), which we called PmcY juxtamembrane (PmcYj), to GST. The fusion protein was purified and its capacity and specificity to bind transition metals analyzed. For this, five molar excess of different metals were incubated with the fusion protein (Fig. 2B-C) and after unbound metal was separated by exclusion chromatography (Fig. 2B) or washed (Fig. 2C), protein was quantified and metal bound to protein detected as described. This strategy allowed us to confirm that Zn^2+^ is the cognate metal for PmcY and that the stoichiometry of binding is 2 Zn^2+^ per PmcY molecule. To identify possible binding sites, we analyzed the Zn^2+^ binding of the single mutants of highly conserved positions with O ligands; *i.e.*, D40A, D47A and D65A. All these mutations lowered the binding stoichiometry to 1, indicating their likely participation in two Zn^2+^ binding sites. A closer look to the distances between O atoms in lateral chains of the PmcY structure, underscores residues D47-D65 as candidates for one Zn^2+^ binding with the archetypical tetrahedral geometry (Fig. 2D).

**Figure 2.**
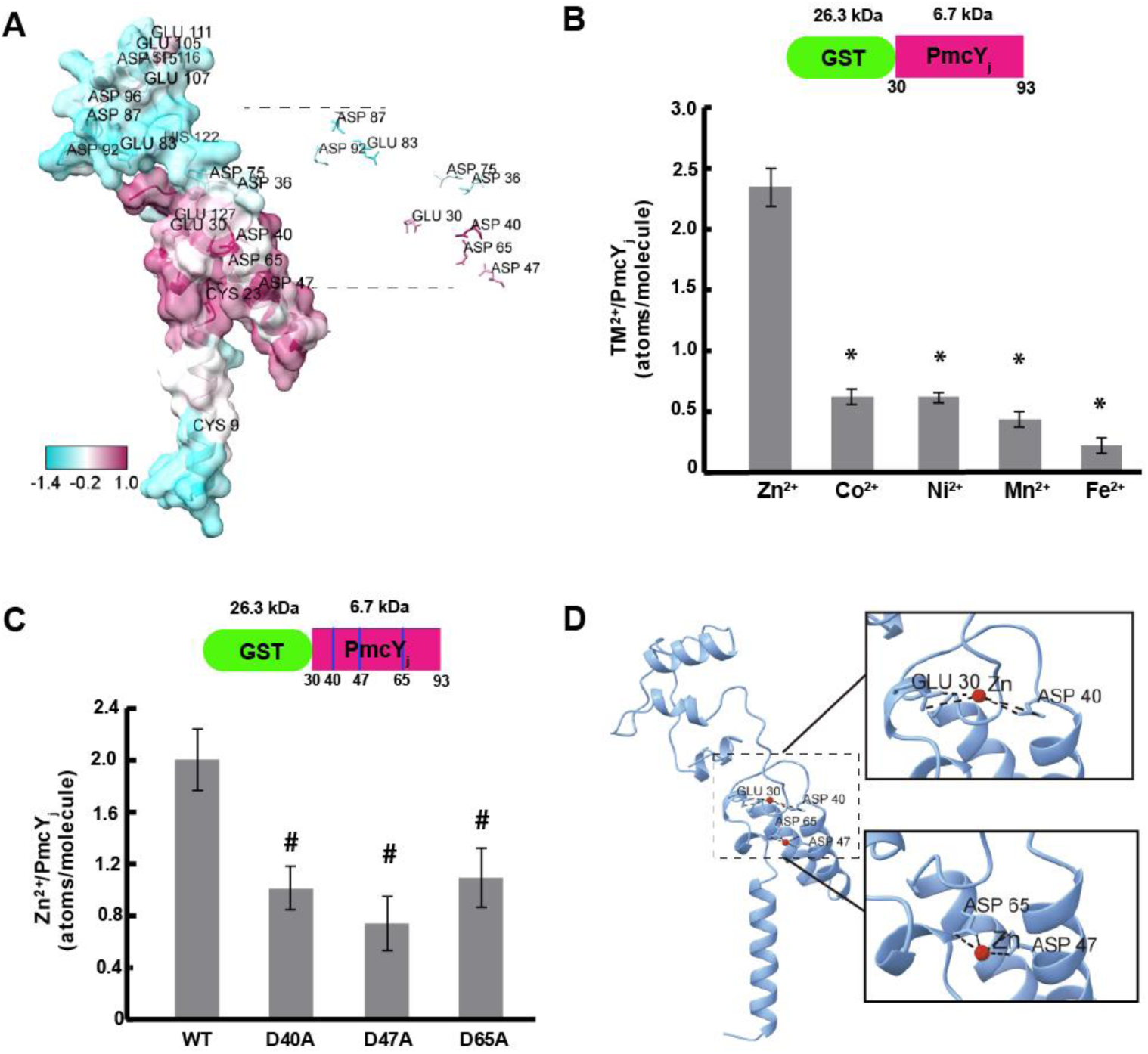
Conserved residues in the juxtamembrane region/domain (PmcYs) have a role in Zn^2+^ binding. (A) Left, conserved amino acids profile depicted in the structure for PmcY/PA3962. Positions colored with maroon indicates high conservation while cyan points to low conservation. Right, lateral chains of putative amino acids with metal binding capacity are shown. (B) Up, representation of the fusion protein GST-PmcYs. Down, metal (TM^2+^) binding capacity and specificity of the fusion product. Metal bound to GST was subtracted in each case and represented 10% of the total metal present in GST-PmcYs. (C) Up, representation of the fusion protein with conserved amino acids positions indicated with blue lines. Down, binding capacity of mutants D40A, D47A and D65A. Notice that Zn^2+^ binding decrease to a half vs. WT. (D) Representation of putative Zn^2+^ binding sites in PmcY. Zn^2+^ atoms are indicated in red and putative bonds by O-ligands with dotted lines. In all cases, data are the mean ± SE of three independent experiments. \**P*<0.01; ^#^, *P*, 0.05 from a two-tailed Student’s *t* test versus Zn^2+^ (B) or WT (C).

### PmcY interacts with the periplasmic Zn^2+^ sensor CzcS

Another property which defines a metallochaperone is the interaction with its target(s) protein(s). This is key in order to promote the ligand transfer reaction, and the metal transfer, between the donor, the metallochaperone, and the acceptor, the target protein. Thus an interaction between PmcY and CzcS could promote Zn^2+^ sensing by means of a ligand transfer reaction between partners. In order to test this, we expressed and purified the periplasmic soluble domain (SD) of CzcS (amino acids 40 to 166, named CzcSSD) as described by Wang et al. (11) with a twin Strep-tag at the N-terminal end. The protein preparation was treated with EDTA to eliminate Zn^2+^ carried-over during purification, and then incubated with GST-PmcYj, in two conditions, EDTA-treated or loaded with Zn^2+^. After collecting flow through and washes we detected the presence of CzcS in the final resin (Rf) only when GST-PmcY_j_ was loaded with Zn^2+^ (Fig. 3A, black asterisk). In order to assess how Zn^2+^ occupancy affected this interaction, GST-PmcYj mutants, D40A, D47A and D65A were analyzed similarly. The Zn^2+^-dependent interaction was lost only in D47A and D65A mutants. This data supports the hypothesis in which these acidic residues might coordinate Zn^2+^ in one binding site. Mutation D40A did not preclude the interaction in presence of Zn^2+^.

**Figure 3.**
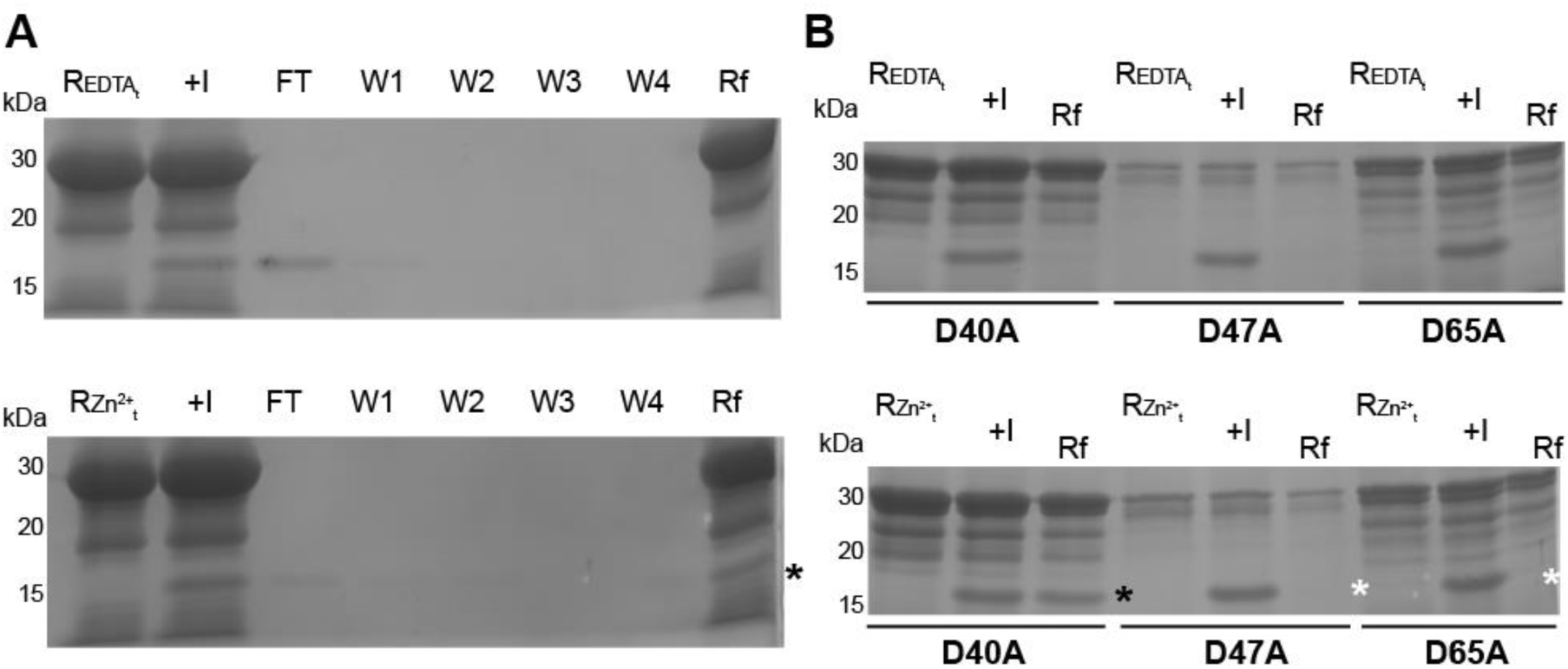
PmcY interacts with CzcS. On-column Zn^2+^-dependent protein-protein interaction assay. (A) Upper and lower panel, 100 mg (200 µl) of resin bound with 10 nmole of GST-PmcYj (PmcY_30-93_) were washed with 10 bed volume of Buffer B (Buffer A + 1 mM EDTA), (Upper panel, lane REDTA_t_), or Buffer C (Buffer A + 0.1 mM Zn^2+^) (Lower panel, lane RZn^2+^_t_) and incubated with 5 nmole CzcS_40-166_ (lane +I). The flow through was collected (FT) and the resin washed 4X with 2 bed volumes of Buffer A (lanes W1-4). Finally, the resin was resuspended at the initial volume (lane Rf) and 25 µl of each fraction analyzed by SDS-PAGE 12% and CBB staining. Note that PmcY-CzcS interaction requires Zn^2+^ bound to PmcY (black asterisk). (B) Protein interaction analyses of PmcY mutants. EDTA treated (upper panel) or Zn^2+^-loaded (lower panel) GST-PmcY mutants (5 nmole D40A, 1 nmole D47A, 2.5 nmole D65A) were incubated with 5 nmole CzcS. Twenty five µl of fractions from resin treated (lanes REDTA_t_ and RZn^2+^_t_), incubated with CzcS (lanes +I) and final (lanes Rf) samples were analyzed as in (A). Black asterisk denotes the Zn^2+^-dependent interaction and white, the lack of it.

**Figure 4.**
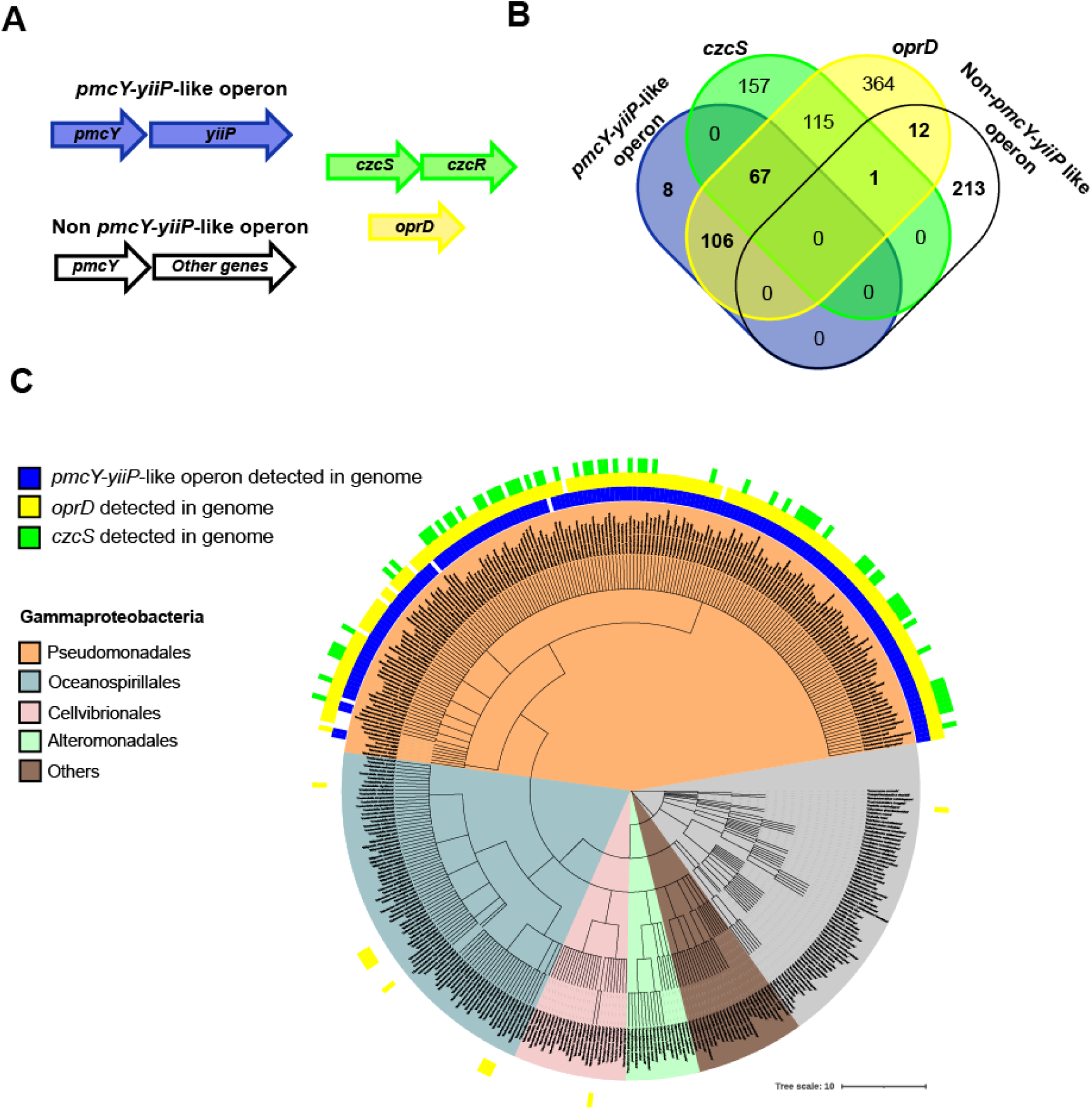
PmcY-YiiP-like operons coevolved with OprD and likely CzcS in Pseudomonadales. Bioinformatic analyses. (A) Two operon configurations involving PmcY homologs were searched in complete bacterial genomes by multigene Blast using protein sequences; *pmcY-yiiP*-like operon (blue) and non-*pmcY-yiiP*-like operon (white). Parallel searches by RBH were done with protein sequences coded by *czcS* (green) and *oprD* (yellow). (B) Venn diagram showing the distribution of organisms with detected CzcS (green) and OprD (yellow) between the two configurations of PmcY (*pmcY-yiiP*-like operon, in blue vs. *non-pmcY-yiiP*-like operon, in yellow). Note the lack of association between *non-pmcY-yiiP*-like operon organisms (226) with CzcS and OprD (1 and 13 organisms among 226, respectively). (C) Phylogenetic analysis of the organisms with PmcY homolog detected in the genome. Most species with *pmcY-yiiP*-like operon (blue strip) belongs to the order of Pseudomonadales (ocher). Coevolution of OprD and CzcS is depicted with yellow and green strips labels, respectively.

### PmcY-YiiP operon coevolved with OprD and CzcS in Pseuodomonadales

The data presented suggest that YiiP, PmcY, CzcS/CzcR participate in a sensing mechanism coupling Zn^2+^ efflux via YiiP, to decrease imipenem suceptibility (as *oprD* expression is downregulated). If this mechanism is conserved among gram-negative bacteria, we hypothesized that PmcY homologs found in other organisms would present the same operon architecture as in *P. aeruginosa* MPAO1. A multigene BLAST search did not support this. However, organisms presenting the operon architecture were mostly of the Pseudomonadales order. Most importantly, contrasted with organisms with non-*pmcY*-*yiiP* operon detected, *oprD* and *czcS* were mostly distributed in organisms with the operon *pmcY-yiiP*; *i.e.*, the Pseudomonadales order. The lack of CzcS in a high number of members in this clade is surprising. However, the periplasmic Cu^+^ sensor CopS has also been proposed to downregulate *oprD* transcription (29). Altogether these data support a likely coevolution of these genes. We discuss in the next section what could be the selection pressure of the coevolution and the physiological role in a context of Zn^2+^ bioavailability.

## Discussion

The association between the Zn^2+^ induced repression of *oprD* transcription, and the Zn^2+^ induced resistance to imipenem is long recognized (30). Moreover, several studies have used this response to dissect important molecular and mechanistic aspects of the CzcS-CzcR signaling pathway (11, 31) in the periplasmic space. Here we present evidence pointing to the participation of YiiP and PmcY in periplasmic Zn^2+^-sensing mechanisms. In particular, we propose that both players affect *oprD* transcription, and likely other genes, by promoting Zn^2+^-sensing; *i.e.*, the metalation of the Zn^2+^ sensor CzcS. We provide evidences supporting our model, showing that PmcY binds Zn^2+^, with high specificity and with a molar stoichiometry of 2 Zn^2+^ per molecule. Metal coordination is given by mean of conserved Asp residues. Phenotypical analyses showed that different to WT and YiiP mutant, the mutant strain of PmcY lacks the Zn^2+^-dependent response to imipenem susceptibility likely by a lack or reduced Zn^2+^-sensing capacity of CzcS. These results lead us to propose PmcY as a Zn^2+^-metalochaperone.

Many of the players involved in orchestrating Zn^2+^ homeostasis in *P. aeruginosa* have been described (32). High Zn^2+^ bioavailability is sensed by periplasmic and cytosolic sensors, CzcS and CadR, respectively which lead to transcription of high capacity Zn^2+^ transport systems, such as HME-RND and PIB-type ATPases (9, 10). The integration of these compartmentalized sensing mechanisms can be interpreted considering electrochemical Zn^2+^ gradient. As Zn^2+^ entries and accumulates first in the periplasm transcription of HME-RND are first expected. However, free Zn^2+^ is almost negligible in cells (33) and a role for cytosolic Zn^2+^ exporters have been proposed in periplasmic Zn^2+^ sensing mechanisms (10). This is important as it suggests the participation of Zn^2+^-transport systems allowing cross-talk between cellular compartments to promote intracellular Zn^2+^ exit even when extracellular Zn^2+^ could turn into lower levels.

Could a similar scenario, involving Zn^2+^ transport systems, participate in cross-talk between compartments when Zn^2+^ bioavailability is low? Considering the risk of losing Zn^2+^ in such scenario, the process should be accompanied of a “favourable” fitness cost. We proposed that Zn^2+^-export by YiiP is linked to Zn^2+^-sensing by CzcS with the final outcome of delaying *oprD* expression, which role allows entry of nutrients, but also xenobiotic molecules such as antibiotics. Thus this hypersensitivity to sense intracellular Zn^2+^, when scarce but still available, would be beneficial for this opportunistic pathogen as *oprD* induction is still repressed.

HME-RND also is part of the CzcS-CzcR regulon (31) and Zn^2+^-binding to CzcS upregulates the transcripts levels. Even when transcription could be triggered by YiiP-PmcY activity we speculate that low Zn^2+^ availability is not sufficient to promote HME-RND activity as this transport system would have similar biochemical properties to CadA; i.e.: high transport capacity but low metal affinity. Incidental Zn^2+^-entry via OprD has been suggested (34). Although counterintuitive, the up-regulation of HME-RND by CzcS activation, would prepare the cell for a potential Zn^2+^ overflow in the periplasm with minimal basal activity of HME-RND until Zn^2+^ bioavailability increases. However, evidence to support this is still missing and the lack of Zn^2+^ resistance shown by OprD mutants contradicts this role (31).

Lastly, and considering our working model, although it is tempting to hypothesize that a Zn^2+^-basic amino acid abduct could be the physiological substrate transported by OprD, the metal-chelating molecule EDTA was not able to outcompete arginine/lysine transport by OprD in *in vitro* studies (5). Nonetheless, considering the Zn^2+^ binding capacity of several xenobiotics, including antibiotics, and the evident lack of specificity of *oprD* for carboxylated substrates, its overexpression could be viewed as a “razors-edge”-like situation promoting Zn^2+^ uptake linked to the xenobiotc molecule carrying the ion. Thus, braking the system could be beneficial until Zn^2+^ reservoirs are downsized to the limits of a “go-hunting” for survival condition.

## Author contributions

Conceived and designed the experiments: D.R., P.M. Analyzed the data: D.R., P.M., T.M., M.E.C.

Contributed reagents/samples/analysis/tools: D.R., M.E.C., M.O.-F., T.P.-B. Wrote or edited the manuscript: D.R., P.M.

All authors read and approved the final manuscript.

## Acknowledgments

We thank Camila Semeniuk, Dr. Laura Montroull and Dr. Andrea Pellegrini for their invaluable help during this work.

## Abbreviations

TM: transition metal
CDF: cation diffusion facilitator
IPTG: Isopropyl β-D-1-thiogalactopyranoside
PAR: 4-(2-Pyridylazo) resorcinol
HME-RND: heavy metal exporter of the resistance-nodulation-division family.

